# Integrated analysis of patterns of DNA breaks reveals break formation mechanisms and their population distribution during replication stress

**DOI:** 10.1101/171439

**Authors:** Yingjie Zhu, Anna Biernacka, Benjamin Pardo, Romain Forey, Norbert Dojer, Raziyeh Yousefi, Jules Nde, Bernard Fongang, Abhishek Mitra, Ji Li, Magdalena Skrzypczak, Andrzej Kudlicki, Philippe Pasero, Krzysztof Ginalski, Maga Rowicka

## Abstract

DNA double-strand breaks (DSBs) can be detected by label-based sequencing or pulsed-field gel electrophoresis (PFGE). Sequencing yields population-average DSB frequencies genome-wide, while PFGE reveals percentages of broken chromosomes. We constructed a mathematical framework to combine advantages of both: high-resolution DSB locations and their population distribution. We also use sequencing read patterns to identify replication-induced DSBs and active replication origins. We describe changes in spatiotemporal replication program upon hydroxyurea-induced replication stress. We found that one-ended DSBs, resulting from collapsed replication forks, are population-representative, while majority of two-ended DSBs (79-100%) are not. To study replication fork collapse, we used strains lacking the checkpoint protein Mec1 and the endonuclease Mus81 and quantified that 19% and 13% of hydroxyurea-induced one-ended DSBs are Mec1-and Mus81-dependent, respectively. We also clarified that Mus81-induced one-ended DSBs are Mec1-dependent.

## Introduction

DNA double-strand breaks (DSBs) are the most deleterious form of DNA damage. DSBs can either arise spontaneously, e.g. during DNA replication or be induced, e.g. by ionizing radiation or chemicals, including many chemotherapy drugs (1). DSBs are also induced during genome editing by the CRISPR/Cas system (2). A better understanding of mechanisms of DSB formation is therefore of great interest and crucial to effective prevention and treatment of the associated diseases, including cancer. Nevertheless, the quantitative knowledge of how genomic DNA breaks in response to various stressors is still lacking. Starting with our BLESS method in 2013 (*3*), several methods for direct DSB-labeling genome-wide have been developed (*4-6*), including i-BLESS (7) used here. These dramatic improvements in DSB detection techniques yielded only modest progress in understanding mechanisms of DSB formation, due to the challenging nature of DSB sequencing data and lack of computational approaches for its more advanced analysis. DSBs are very rare events, even under DSB induction, since cells can only tolerate few DNA breaks. Therefore, even a fraction of a percent of e.g. senescent or apoptotic cells exhibiting massive DNA fragmentation can obscure the DSB sequencing data.

Here, we address this challenge by proposing a computational framework to analyze subpopulations of cells with different DSB patterns (**Fig. 1a**). The method relies on DSB sequencing and pulsed-field gel electrophoresis (PFGE) (**Fig. 1b**) as main input data, which we integrate computationally. The PFGE is a method of choice to separate large DNA fragments of up to several megabases and to monitor the integrity of chromosomes. This technique is less sensitive than sequencing in detecting the presence of DSBs, but, unlike DSB sequencing, it cannot be obscured by a small subpopulation of highly-damaged cells (**Fig. 1c**). Our newly introduced quantitative DSB-sequencing (qDSB-Seq) (*8*) method allows us to quantify the number of DSBs per cell (**Fig. 1a**), which is key to understanding their physiological relevance. To further study DSB formation, we classify DSBs into patterns related to their generation mechanisms. Such classification allows to quantify the contributions of different sources of DNA damage in a given condition.

**Figure 1.**
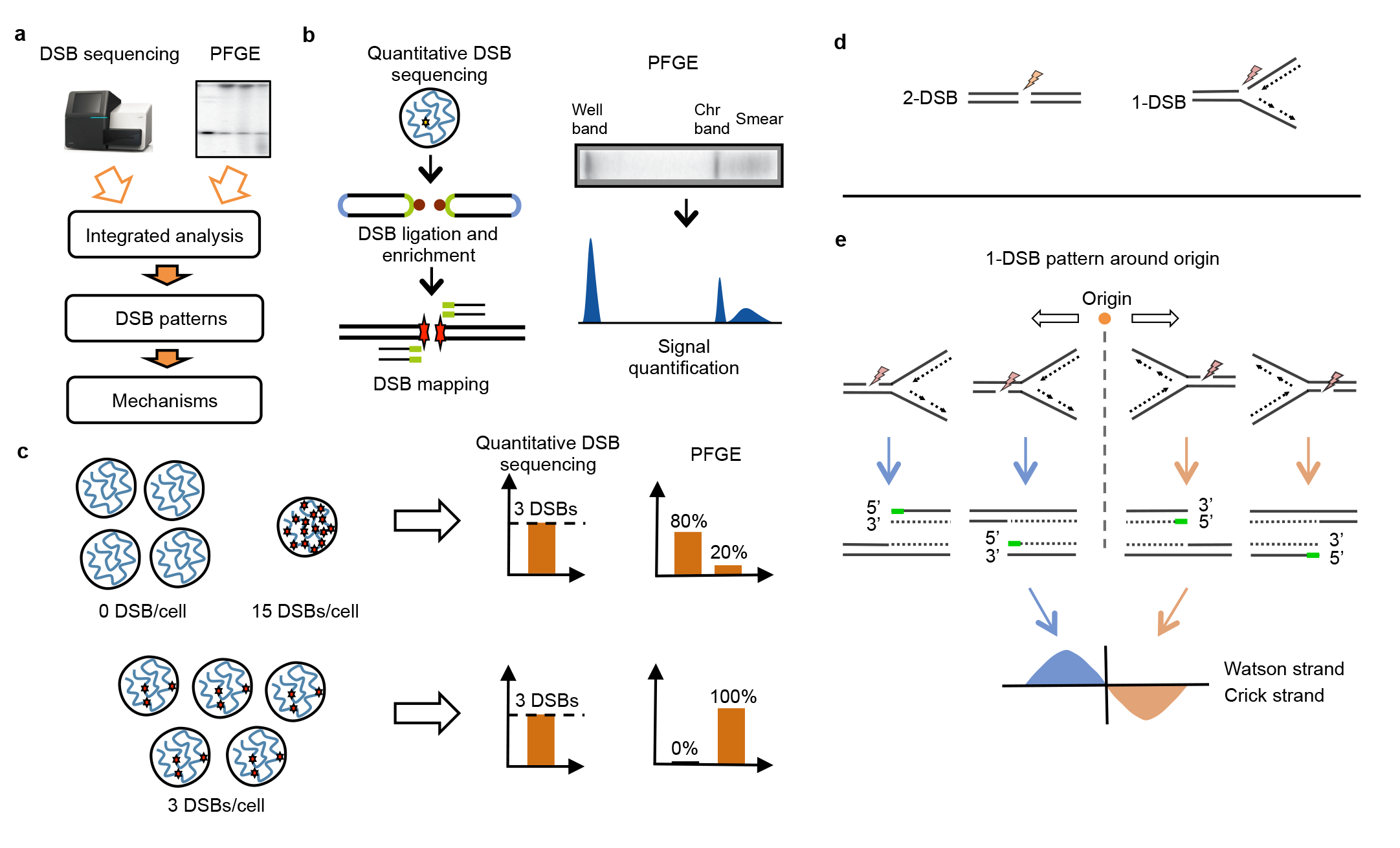
Integrative analysis of DSB formation. **(a) Overall approach.** Different data sources are integrated computationally, DSB formation mechanisms and their prevalence in population are inferred. **(b) Main data sources:** DSB-sequencing and pulsed-field gel electrophoresis. **(c) Differences between DSB-sequencing and PFGE.** DSB-sequencing reports an average number of DSBs in population, so the result may not reflect typical DSB number in population, even if small subpopulation of cells with high number of breaks is present (e.g. dead cells). PFGE detects lengths of DNA fragments, so it does not have a problem between distinguishing if breaks occur in a small population of cells or are representative of majority of population. **(d) 2-ended and 1-ended DSBs. (e) Expected DSB pattern resulting from fork collapse.** The double-strand breaks are generated when replication fork passes through the single-strand breaks (nicks). The collapse of forks travelling to left and right will result in reads mapped to the + and - strands, respectively. The expected total pattern is shown below.

We show how our framework can be applied to studying a complex repertoire of one-ended and two-ended DSBs induced by replication stress. One-ended DSBs (1-DSBs, **Fig. 1d**), are a unique type of DSBs resulting from collapsed or reversed replication forks, while all other DSBs are two-ended (2-DSBs). 1-DSBs manifest as an asymmetric sequencing read pattern (**Fig. 1e**), which can be utilized to detect active replication origins. Learning DSB patterns related to specific DSB-inducing mechanisms allows classifying breaks according to mechanisms of their formation and quantify sources of DNA damage.

## Results

To evaluate DSB patterns caused by replication stress, we arrested haploid yeast cells in G1 and released them synchronously into S phase in the presence of hydroxyurea (HU), which slows replication by depleting dNTPs. To study the mechanisms of stabilization and repair of stalled forks, we used strains lacking the checkpoint protein Mec1 and the endonuclease Mus81, which was proposed to cleave stalled forks (**Fig. 2a**). i-BLESS was used for DSB labeling (7), followed by qDSB-Seq to quantify DSBs (*8*). To better understand the DNA damage induced by replication stress, we analyzed the resulting DSB read patterns and classified DSBs according to their creation mechanisms.

**Figure 2.**
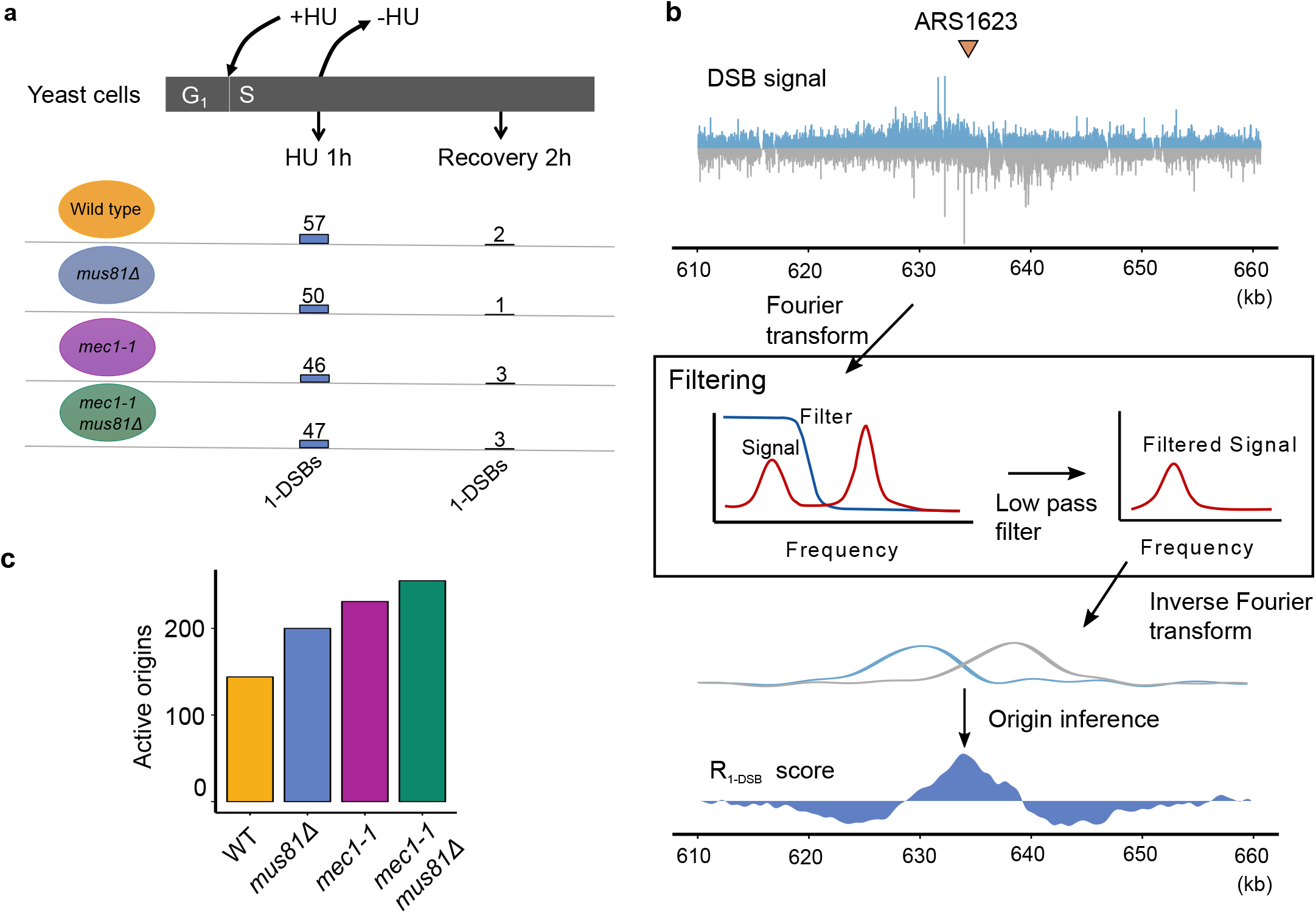
Quantification of 1-DSBs in replication stress and release conditions and active origin inference from 1-DSB signal. **(a) Experimental conditions and results of 1-DSB quantification.** Yeast cells were synchronously released from *α*-factor G_1_ phase arrest, treated with HU for 1h, and resuspended in fresh medium to recover from HU. Quantification of 1-DSBs is explained below. **(b) Active origin inference after noise filtering.** DSB-sequencing data were transformed to frequency signal using Fourier transform, then low pass filter and inverse Fourier transform was executed to remove high frequency signal. R_1-D_sb score was calculated to identify origin and range of replication. **(c) Numbers of origins detected in HU-treated samples.**

### Identification of replication-dependent 1-DSBs

For an individual DSB-sequencing read, it is impossible to distinguish whether it originates from a 1-DSB or 2-DSB. Therefore, to identify true 1-DSBs, originating from fork collapse, we first determined active origins and their replicated vicinity (fork range). Briefly, we mapped the i-BLESS-labeled (7) and sequenced DSB reads separately into the + and - strands of the yeast genome and scored the DSB distribution on its resemblance to the origin pattern (**Fig. 1e, Methods**). To facilitate pattern detection, we removed the high-frequency noise (**Fig. 2b**), e.g. related to nucleosome spacing (**Fig. S1**), while preserving lower-frequency components related to replication-dependent DSBs. This computational method gives results very similar to optimized experimental DSB detection protocol (**Fig. S2**). In thus filtered data (**Fig. 2b**), we used a sliding window to find regions with the highest 1-DSB score (R_1-DSB_) (**Methods Eq. 2-4**). Finally, the hypergeometric test was used to estimate the significance of the R_1-DSB_ scores and thus select predicted origins (**Fig. S3**).

This method yields highly precise and sensitive origin predictions (**Table S1, Fig. S4**). For example, for HU-treated *mec1-1 mus81Δ* cells, we detected 285 origins (at 95% precision, 85% sensitivity) or 255 origins (at 100% precision, 83% sensitivity). Hereafter, 100% precision origin predictions are used (**Fig. 2c**).

Prediction of active replication origins allowed us to distinguish between breaks resulting directly from replication fork collapse and others. We estimated “fork range”, that is the maximal region detected as replicated by a given origin, as a length of an interval exhibiting a statistically significant preference for 1-DSBs caused by collapsed forks from that origin (**Fig. 1e, Methods**). Only 1-DSBs detected in these fork ranges were accepted and only those for which a corresponding read on the other strand, which could originate from a putative 2-DSB, could not be found in the vicinity. Thus, we classified DSBs into 1-DSBs and 2-DSBs.

### Inferring DSB distribution across cell population

To interpret the calculated average numbers of DSBs per cell, it is crucial to know if they are representative of the majority of cells in the population (**Fig. 1c**, top vs. bottom). To infer the distribution of DSBs in a cell population, we utilized pulsed-field gel electrophoresis (PFGE), which allows separation of DNA fragments from 50 kb to several megabases, the size of the largest budding yeast chromosomes. During PFGE, replicating chromosomes cannot enter the gel due to replication intermediates and therefore remain in the well (**Fig. 1b**, and **Fig. 3a**, **structures 3-4**). Non-replicating chromosomes enter the gel and form a distinct band, corresponding to intact chromosomes, whereas the fragmented ones are detected as a smear (**Fig. 1b**). We transferred gels on nylon membranes to detect the chromosome III with radiolabeled probes (Southern blot) for HU-arrested (HU) cells and those recovered from the treatment (Recovery, **Fig. 2a**). Each broken, non-replicating chromosome III would contribute equally to the smear signal, irrespective of the number of 2-DSBs. Therefore, the PFGE data is not affected by a small fraction of cells with many DSBs (**Fig. 1c**) and thus perfectly complements DSB-sequencing data by providing information on typical distribution of DSBs in cell population.

**Figure 3.**
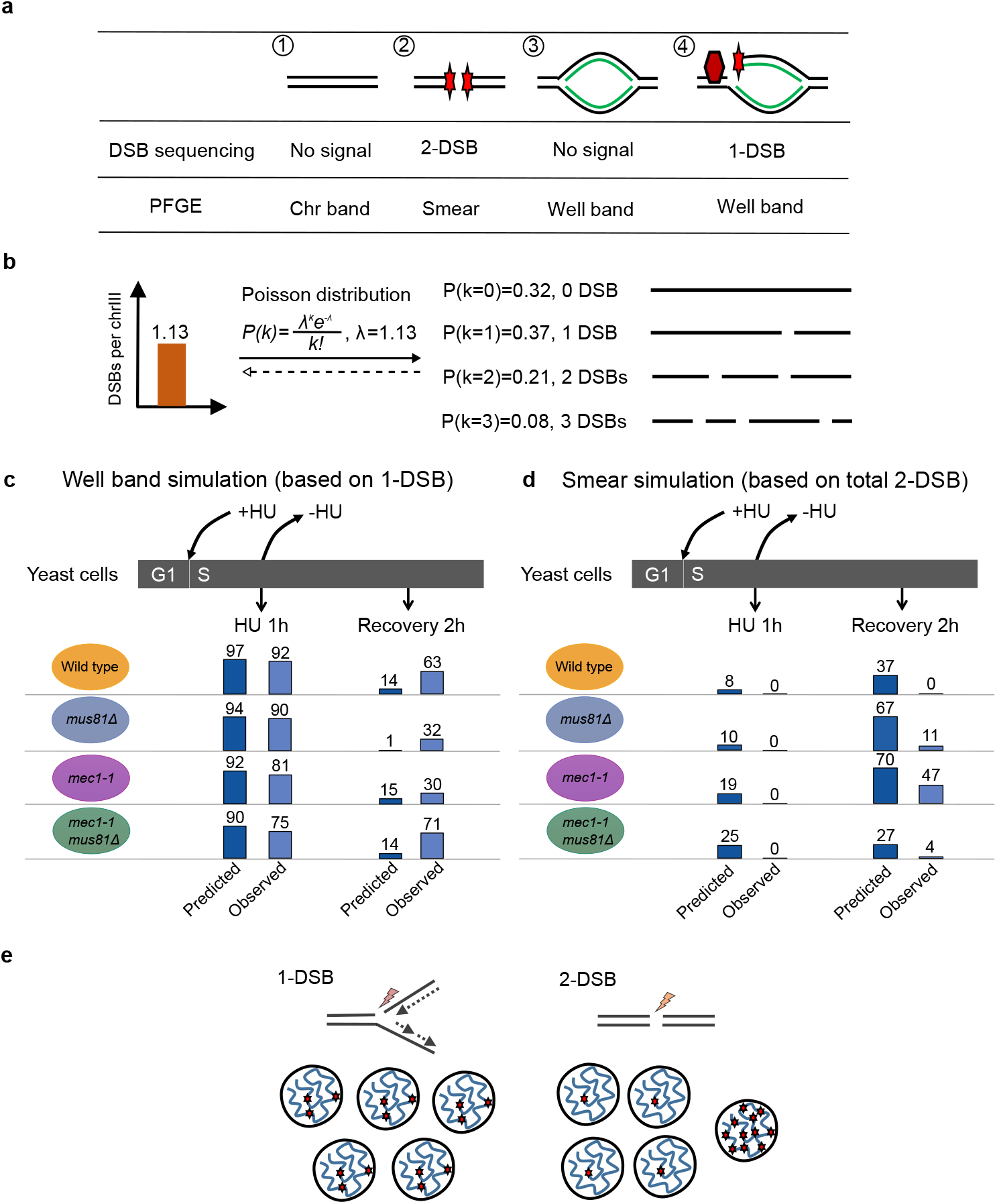
Comparison of qDSB-Seq and PFGE results. **(a)** Results of DSB sequencing and PFGE for different chromosomal fragments: (1) Unbroken linear DNA; (2) Linear DNA with 2-DSB; (3) Replicated, but unbroken DNA; (4) Replicated DNA with 1-DSB. **(b)** An example of use of the Poisson distribution to simulate population-wide DSB distributions using the measured number per chromosome III (here 1.13). **(c)-(d)** Comparison of predicted (Poisson model) and observed (PFGE) well band and smear band on chrIII: **(c)** Well band simulation based on qDSB-seq 1-DSB data; **(d)** Smear simulation based on qDSB-seq fork 2-DSBs. **(e)** Cell population of 1-DSBs and 2-DSBs. Small subpopulation with high number of 2-DSBs cannot be detected by PFGE.

DSB-sequencing gives the average number of DSBs per cell and PFGE data provide percentage of replicating, non-replicating intact and non-replicating broken chromosomes III (well-band, chromosome-band and smear signal, respectively, **Fig. 1b**). To compare DSB-sequencing and PFGE results we needed to model one data type using the other. First, we modeled 1-DSB distribution in the genome using the Poisson distribution (**Fig. 3b**), i.e. assuming that 1-DSBs occur independently of each other. Both 1-DSBs and intact replication intermediates contribute to the well-band signal, so the percentage of DNA in the well band predicted from 1-DSBs alone should be not greater than the observed one. Nevertheless, our predictions are 4% to 20% higher than observed values in HU samples (**Fig. 3c**). This small difference may reflect the fact that not all replication intermediates remain trapped in the well during the PFGE.

Moreover, 1-DSBs can be only formed from branched DNA structures, such as collapsed replication forks (**Fig. 1d**). Based on our computer simulations of DNA replication (*9*) and using FACS data to estimate replication progress, we calculated that on average there are 168 replication forks in wild-type cells. qDSB-Seq estimated 57 1-DSBs per cell, meaning that on average 34% of all replication forks collapsed to 1-DSBs. Even in case of forks forming 1-DSB preferentially in a subpopulation of cells, such subpopulation would be major, at least one-third of all. Since no data we gathered suggest such preferences exist, we conclude that in HU samples 1-DSBs occur in virtually every replicating chromosome III. Similar calculations for recovery samples show that few (3-50%, **Fig. 3c**) replicating chromosomes III exhibit 1-DSBs, suggesting an efficient 1-DSBs repair is occurring during 2-hour recovery from the HU treatment. These data also indicate that 1-DSBs are occurring frequently during fork stopping.

Next, we employed the Poisson simulations to predict PFGE smear signal. Predictions were not consistent with observations (**Fig. 3d**) and indicated that 2-DSBs are highly preferred in already broken chromosomes. We also estimated average 2-DSB numbers per chromosome III in majority of the population, using PFGE data to predict corresponding average DSB levels expected to be measured by qDSB-Seq (**Fig. 3b**). We concluded that virtually all 2-DSBs detected by sequencing in HU samples originated from a cell subpopulation too small to be detected by PFGE (**Fig. 3e**), but having so many 2-DSBs that it overpowers the sequencing data (**Fig. 1c**). For example, a single yeast cell with DNA fragmented into units corresponding to single nucleosomes would give rise to ~70,871 DSBs (*10*). Therefore, 0.3% of such cells would contribute 240 DSBs/cell to population mean. Another indication of the presence of small population of cells with highly fragmented DNA is that we detect much more breaks within several hundred base pair distance from each other than expected by chance (**Table S2**). Such breaks are distributed evenly within the genome and their level is changing with conditions, indicating that they may originate from highly damaged cells, which percentage is condition-dependent. We conclude that such a small cell subpopulation, undetectable by PFGE, is a source of 100% of 2-DSBs in HU samples.

In recovery samples, similar reasoning indicates that small population of highly-damaged cells gives rise to 100% of 2-DSBs for wild-type, 96% for *mus81Δ*, 95% for *mec1-1 mus81Δ* and 79% for *mec1-1* (**Fig. S6**). The remaining 2-DSBs are confirmed by PFGE. We conclude that they do not result directly from fork collapse, since the 2-DSB distribution does not reflect stalled fork positions, but sequencing read density (**Fig. S8**), suggesting they are random. Moreover, FACS data (**Fig. S7**) show that in *mec1-1* and in *mec1-1 mus81Δ* cells replication cannot be completed, unlike in other samples. Taken together, these data suggest that in *mec1-1* cells problems with completing replication lead to 2-DSBs. Percentage of broken chromosomes detected by PFGE in *mec1-1 mus81Δ* recovery sample is much lower than in *mec1-1* cells. It may be related to Mec1 protecting fork from Mus81-mediated cleavage, with removal of this protection causing PFGE-detected breaks in *mec1-1* cells during recovery.

Since 1-DSBs detected by sequencing are population-representative and 2-DSBs typically are not, in the subsequent analysis we focus on 1-DSBs.

### Role of Mec1 in origin firing

The Mec1 protein was proposed to inhibit firing of late origins during replication stress (*11-13*). Indeed, we detected 60 active late origins in *mec1-1* cells and only 7 in wild-type cells (**Fig. 4a**). We also observed increased number of overall origins activated under HU treatment in *mec1-1*, 231 vs. 144 for wild-type (**Fig. 2c**). On the other hand, number of 1-DSBs per cell directly originating from fork collapse, estimated conservatively as described above, varied less: 57 for wild-type cells and 46 for *mec1-1*. Assuming that stopped forks have the same (or not higher) chances of progressing to 1-DSB in wild-type compared to *mec1-1* cells, this result suggests that even though the number of fired origins increases in Mec1-deficient cells, their efficiencies (i.e. percentage of cells in which a given origin is fired) decreases about two-fold (**Fig. 4b**). This result supports the idea that Mec1 promotes origin firing (*14*).

**Figure 4.**
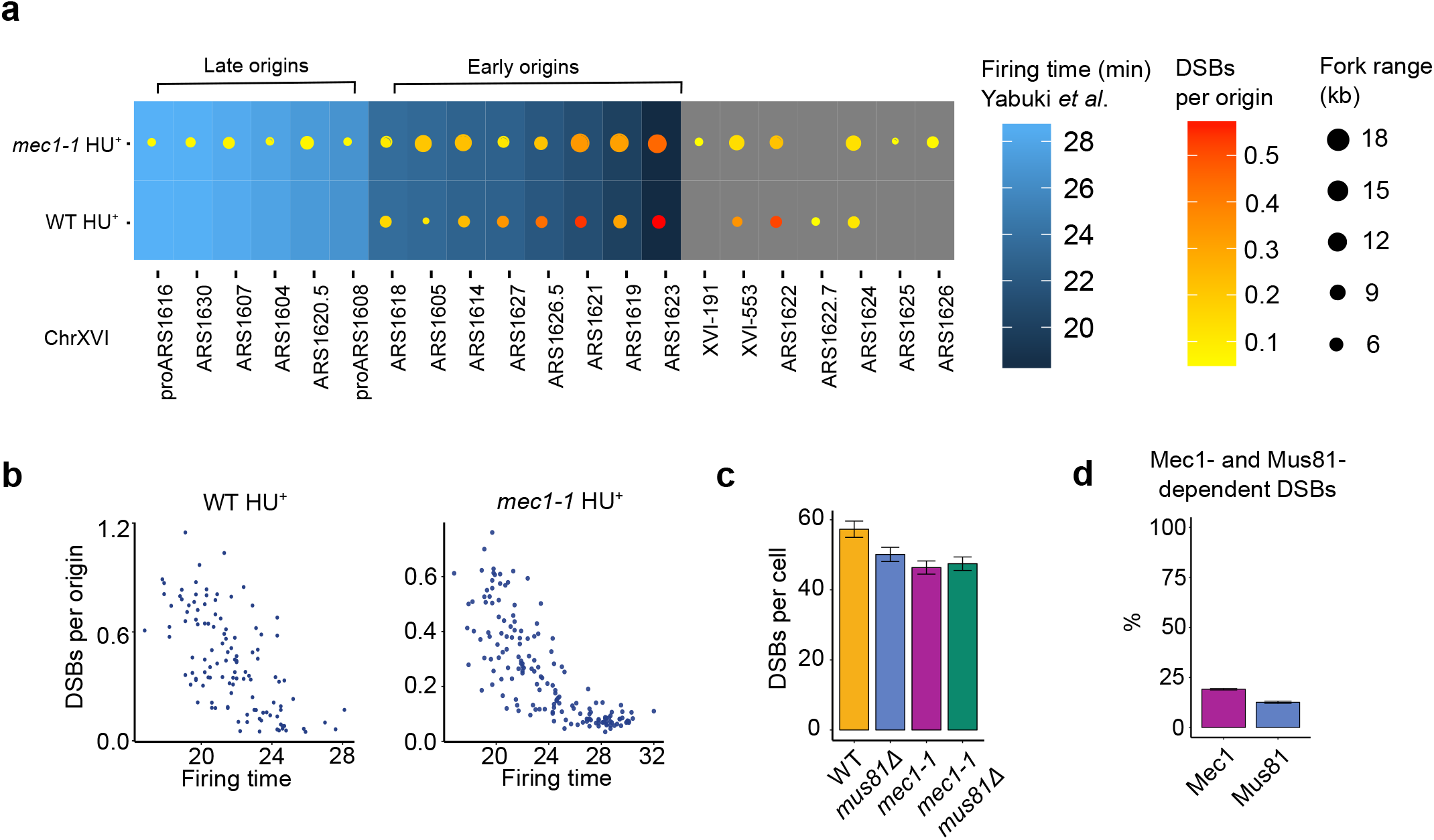
The roles of Mec1 and Mus81 in response to replication stress. **(a)** Visualization of fork ranges and 1-DSBs for selected origins. **(b)** Correlation between replication-dependent 1-DSBs and firing time. Lower 1-DSB level was observed in *mec1-1* mutant cells compared to wild-type. **(c)** Quantification of 1-DSBs per cell. **(d)** Mec1-and Mus81-dependent DSBs in HU-treated wild-type cells.

### Role of Mec1 and Mus81 in resolving replication stress

Mus81 is a catalytic component of a eukaryotic structure-selective endonuclease and was proposed to cleave stalled replication forks, nevertheless its role in resolving replication stress remains incompletely understood (*15-17*). In physiological conditions, replication-induced DSBs are resolved quickly by orchestrated actions of DNA damage response proteins, such as Mec1. We observed that number of 1-DSBs is decreased by 13% upon Mus81 deletion and by 19% in Mec1-deficient cells (**Fig. 4c**). Moreover, in Mec1-deficient cells, deletion of Mus81 does not impact number of 1-DSBs. Taken together, we show that the vast majority of the 1-DSBs in HU-treated wild-type cells are Mus81 - independent, meaning that they are not caused by Mus81 cleavage of stalled or reversed forks. Almost as many HU-induced 1-DSBs, 81%, does not depend on Mec1. Interestingly, in Mec1-deficient cells, deletion of Mus81 does not change the number of breaks, indicating that cleaving of stalled or reversed forks by Mus81 is Mec1-dependent (**Fig. 4d**).

## Discussion

Several methods for direct DSB-labeling genome-wide that have been developed recently allowing DSB detection with unprecedented single nucleotide resolution. Since DSBs are rare in healthy cells, DSB-sequencing data from such cells can be easily overshadowed by a small fraction of the cell population with a substantial DNA damage. This challenge is unique to DSB sequencing, as noise orders of magnitude higher than signal of interest is not normally encountered in any other genomic or omics data.

Therefore, to draw correct conclusions from DSB-sequencing data, it is crucial to consider not only directly measured average DSB numbers per cell, but also how they are distributed in the population. The best solution is to complement DSB sequencing with additional experiments allowing estimating the distribution of DSB counts across the population of cells. Here, we used pulse-field gel electrophoresis and provided conceptual and mathematical framework for integrating these data. Such analysis is crucial for understanding the physiological relevance of measured data and origins of the observed breaks.

In our case, 1-ended breaks that we can directly link to fork collapse tend to be representative of the major population. This is likely a typical situation, if care is exercised to identify only genuine 1-DSBs. Due to their unique nature, originating from collapsed replication forks, the maximal theoretically achievable number of 1-DSBs is twice the number of replication origins, corresponding to 1-DSBs every ~25kb in yeast. On the other hand, 2-DSBs can be caused by any mechanism, such as irradiation, but also replication stress (e.g. breaking of non-replicated DNA). Some mechanisms, such as radiation, may affect all cells to a similar extent in terms of number of DSBs caused. On the other hand, certain other phenomena, such as replication stress, may lead to a low number of 2-DSBs in majority of cells, but in few cells unable to resolve replication stress high number of 2-DSBs may be accumulated, like in the examples discussed above. 2-DSBs from such cells have high potential to obscure the data. For example, in cells undergoing apoptosis, DNA is fragmented into nucleosome-sized units, thus creating several orders of magnitude more breaks than in a typical cell in a population. Frequency of adjacent DSBs also indicates that 2-DSBs are highly enriched in some cells. The observed high numbers of 2-DSBs are not an artifact of our DSB-labeling method, since they are present only in S phase samples (*18*).

This example shows that careful analysis is needed to interpret DSB-sequencing data. Simplistic solutions such as rejecting all 2-DSBs would not be correct, as we showed that some of them are population representative. We also showed that analysis of the genome-wide distribution of DSB patterns allows drawing inference and state hypotheses about mechanisms of their formation. In our case, distribution of 2-DSBs in the fork regions is not consistent with them being caused by fork re-start and indicates random genome fragmentation.

Moreover, analysis of DSBs originating from the collapsed forks gives also insights into the dynamics of unperturbed replication forks and cell replication program. We developed methods for highly precise detection of the active replication origins using DSB-sequencing data, which can be interesting on its own. For example, we discovered that in addition to its known role in inhibiting late origins, Mec1 seems to increase significantly, on average twice, the origin efficiencies.

We also show that careful DSB quantification and classification of breaks according to their mechanisms allows new mechanistic insights. Here, we clarified the role of Mus81 in replication stress, showing that Mus81-mediated breaks are Mec1-dependent. We also showed that Mus81 is responsible for 13% of DSBs induced by HU and that 19% of such DSBs are Mec1-dependent (**Fig. 4d**). That is an important finding, showing that the vast majority of HU-induced 1-DSBs are not caused by cleaving of stalled or reversed forks by Mus81.

In conclusion, we showed how even typically undetectable population heterogeneity can severely contaminate DSB-sequencing data and how this problem can be solved by a computational integration of complementary data types, here by including PFGE in the analysis. Due to DSB-sequencing data susceptibility to noise from few highly damage cells, to ensure correct interpretation of DSB-sequencing data it is critical that distribution of DSBs across cell population will be interrogated.

## Methods

### Strains and growth conditions

Yeast strains used in this study (W303) are listed in **Table S3**. Cells were grown in YPD medium at 25°C until early log phase and were then arrested in G_1_ for 170 min with 8 μg/ml *α*-factor. YBP-275 strain was cultured in YPR medium; galactose was added for 2 h to induce I-SceI cutting. Cells were released from G1 arrest by addition of 75 μg/ml Pronase (Sigma), 200 mM HU (Abcam) was added 20 min before Pronase release followed by 1 h incubation. The cells were subsequently centrifuged and resuspended in fresh medium to recover for 2 h. After collecting, cells were fixated with 2% formaldehyde for 5 min, washed with cold SE buffer (5M NaCl, 500 mM EDTA, pH 7.5) and subjected to DSB labelling.

#### DSB labelling and sequencing

*i*-BLESS DSB labeling was performed as described in (*7*)(*19*). Sequencing libraries for i-BLESS and respective gDNA samples were prepared using commercially available kits ( ThruPLEX DNA-seq Kit (Rubicon Genomics). i-BLESS libraries were prepared without prior fragmentation and further size selection. Quality and quantity of the libraries were assessed on 2100 Bioanalyzer using HS DNA Kit (Agilent), and on Qubit 2.0 Fluorometer using Qubit dsDNA HS Assay Kit (Life Technologies). The libraries were sequenced (2x70 bp or longer) on Illumina HiSeq2500 platform, according to our modified experimental and software protocols for generation of high-quality data from low-diversity samples, such as resulting from the i-BLESS (*19*). Additionally, qDSB-sequencing was performed, either using NotI restriction digestion or I-SceI spike-in, as described in (*8*).

### Quantification of DSBs

We used qDSB-seq method to normalize and quantify DSBs, as described previously (*8*). First, we used YBP-275 strain with Gal-inducible I-SceI endonuclease and I-SceI recognition site introduced at ADH4 locus on chromosome VII. Next, we performed i-BLESS and genomic DNA sequencing. Genomic DNA was used to quantify DSBs by estimating cutting frequency of the enzyme used (*8*).

### Sequencing data analysis

We used *InstantSeq* (*20*) to ensure sequencing data quality before mapping. Next, *InstantSeq* was used to remove i-BLESS proximal and distal barcodes (TCGAGGTAGTA and TCGAGACGACG, respectively). Reads labeled with the proximal barcode, which are directly adjacent to DSBs, were selected and mapped to the yeast S288C genome and the human hg19 genome using bowtie (21) v0.12.2 with the alignments parameters ‘-m1-v1’ (to exclude ambiguous mapping and low-quality reads). To correct single nucleotide variants of yeast strains, we mapped genomic reads of yeast strain used in our experiments to S288C genome using bowtie v0.12.2 allowing 2 mismatches. The reference base pairs were corrected if the variant was present in > 95% cases. The end base pairs of the reads were trimmed using bowtie ‘-3’ parameter. The parameter choice was based on the *InstantSeq* quality report. Hygestat_BLESS v1.2.3 (part of *InstantSeq* software suite (20)) was used to identify genomic regions with a significant difference in normalized read numbers between treatment and control samples.

### Fourier-based low-pass filter

The i-BLESS data contain high frequency quasi-periodic components related to nucleosome spacing and potential sequencing artifacts (apparent in the power spectrum of the read count distribution, **Fig. S1**). To remove the spurious signal from further analysis, we implemented and applied a low-pass filtering approach. The low-pass filter was performed by multiplying the Fourier transform, *D(f)*, of the read density *d(x)*, by a kernel *k(f)*, defined as a smoothed step function

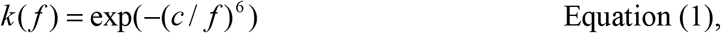

where *f* is frequency and *c* is the cut-off frequency (here we used c=2π/5000 nt^-1^). After filtering in the frequency domain, the data are transformed back to position in the real domain (chromosomal coordinate) by inverse Fourier transform, thus the filtered read density d_filtered_(x) is calculated as the inverse Fourier transform of

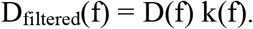

The exponent 6 in the kernel (**Eq. 1**) has been chosen empirically to assure a smooth transition while maintaining an efficient separation of sequencing and nucleosome-related noise from the low-frequency components related to the replication domains. The Fourier transforms were implemented by real-to-complex routines from the FFTW library.

This Fourier-based low-pass filter, which by definition removes high-frequency data, increases signal-to-noise ratio in our DSB-sequencing data. Low-pass filter noise removal method proved very effective, giving results very similar to those obtained by optimizing DSB detection procedure aimed to minimize noise (**Fig. S2**). The low-pass filtering is especially crucial in case of a weaker signal or noisier data (**Fig. S2**, top panel).

### Predicting replication-induced 1-DSBs

We first calculated the numbers of the DSB reads mapped to the left of a candidate origin, mapped to the Watson (WL) and Crick strands (CL), respectively. Analogously, CR and WR were calculated. To eliminate 2-DSBs, we used the difference of read depth between both strands as an estimate of number of 1-DSBs in a region:

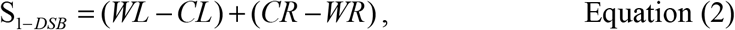

Analogously, the estimated number of 2-DSBs is:

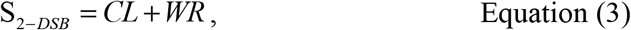

These are conservative estimates; they may lead to underestimating DSBs caused by fork collapse in regions where breaks from forks travelling in both directions are present.

To identify replication origins and fork meeting locations, we defined a ratio for 1-ended DSB (R_1-DSB_), based on the model of replication-induced 1-DSBs (**Fig. 1c**):

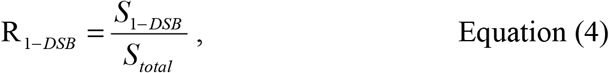

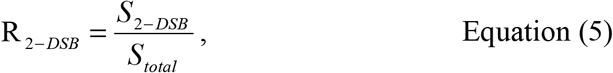

R_1-DSB_ ranges from −1 to 1. R_2-DSB_ ranges from 0 to 1. In the case of an ideal candidate origin with only replication-induced 1-ended DSB, R_1-DSB_ = 1, R_2-DSB_ = 0. In the case of an ideal fork meeting region, R_1-DSB_ = −1, R_2-DSB_ = 1. In the case of an ideal 2-DSB only region, R_1-DSB_ = 0, R_2-DSB_ = 0.5 (**Fig. 2b**).

R_1-DSB_ score encodes a simplified origin model. More specific models can be constructed, but since they would depend on origin firing time and progress of replication at the time when sample was taken. Therefore, we prefer a more general R_1-DSB_ score, which proved very efficient in identifying candidate origins. We used sliding windows ranging from 2 kb to 20 kb to identify candidate origins (local maxima of R_1-DSB_ score). Once candidate origins were identified, the fork range was defined as the longest range (in multiples of 1 kb), where the substantial (>0.5%) enrichment of 1-DSBs was observed in each consecutive 1 kb interval. Then, significance of the enrichment of 1-ended DSB in thus defined fork range was calculated using hypergeometric test, only candidate origins with significant p-values for 1-DSB enrichment were selected as predicted origins.

### Pulsed-field gel electrophoresis

Pulsed-field gel electrophoresis (PFGE) was performed as described previously (*22*). Gel electrophoresis image was analyzed using GelAnalyzer (http://www.gelanalyzer.com/) and ImageJ (https://imagej.nih.gov/ij/index.html). Probes of ARS305, 307, 315 on chrIII were used.

